# MDMA enhances prefrontal plasticity and representational drift during fear extinction

**DOI:** 10.64898/2026.03.06.710094

**Authors:** Nitzan Geva, Sarah J. Jefferson, Emi Krishnamurthy, Tanner L. Anderson, Jocelyne A. Rondeau, Patrick H. Wehrle, Axel F. Rosado, Christopher Pittenger, John H. Krystal, Alfred P. Kaye

## Abstract

Fear extinction requires dynamic updating of cortical representations, yet the neural mechanisms underlying successful extinction remain poorly understood. Some psychoactive substances induce structural plasticity in medial prefrontal cortex (mPFC), possibly underlying their therapeutic potential. Here we investigated whether MDMA, which enhances fear extinction, induces prefrontal structural and functional plasticity, and measured its effects on ensemble representations during extinction. Longitudinal two-photon microscopy revealed that MDMA increased spine density and spinogenesis across prefrontal subregions. Miniscope Ca²⁺ imaging in infralimbic cortex (IL) during fear extinction revealed that IL became more correlated with the suppression of freezing behavior, consistent with a strengthening of its role in extinction. Longitudinal cell registration demonstrated accelerated representational drift across days in MDMA-treated mice; this effect was strongest in a functionally defined subpopulation of neurons that showed suppression of activity to conditioned cues. These findings demonstrate that MDMA facilitates structural and functional neuroplasticity, potentially underlying its enhancement of extinction learning.

## Introduction

Fear extinction requires the dynamic updating of cortical representations, with the medial prefrontal cortex (mPFC) and specifically the infralimbic cortex (IL), playing a central role in both the acquisition and extinction of conditioned fear responses (Milad & Quirk, 2002; Giustino & Maren, 2015; Do-Monte et al., 2015). However, the precise neural mechanisms underlying successful updating of extinction representations remain poorly understood.

Recent work has shown that rapid-acting antidepressants, including ketamine and psilocybin, induce rapid structural plasticity in the mouse medial prefrontal cortex (mPFC), characterized by increased dendritic spine density that may persist for weeks following a single administration (Li 2010; Shao et al., 2021; Olsen, 2018; Jefferson et al., 2023). These structural changes are thought to represent a window of enhanced plasticity which might underlie the therapeutic potential of these drugs for treatment of psychiatric conditions such as depression.

3,4-Methylenedioxymethamphetamine (MDMA) has recently emerged as a breakthrough therapeutic agent in the treatment of post-traumatic stress disorder (PTSD) and has been shown to enhance fear extinction learning in both mice and humans (Young et al., 2015; Young et al., 2017; Vizeli et al., 2022). Additionally, Phase 3 clinical trials have demonstrated significant reductions in symptom severity when psychotherapy is augmented with MDMA (Mitchell et al., 2021; 2023). Despite these initial clinical advances, FDA rejected a new drug application for this treatment and the neural mechanisms underlying MDMA’s potential therapeutic efficacy remain poorly understood. Therefore, we asked whether MDMA, despite having a different pharmacological profile from classical psychedelics, also induces structural neuroplasticity, and whether such changes relate to its therapeutic effects on fear-related learning.

In this study, we used longitudinal two-photon microscopy to track individual dendritic spines in the mPFC over multiple weeks after MDMA administration. We found that MDMA robustly increased spine density in mPFC regions involved in fear acquisition and extinction. Using quantitative proteomics of mPFC synaptosomes, we found that MDMA elevated levels of neurofilament proteins at the synapse. Slice electrophysiology showed that MDMA also enhanced functional plasticity through an increase in spontaneous excitatory postsynaptic current (sEPSC) amplitudes. We then used longitudinal miniscope Ca²⁺ imaging to track ensemble activity in IL during fear extinction and found that MDMA treatment altered the functional dynamics of neural ensembles, reducing the similarity of cue-evoked responses across days.

Together, these findings reveal that MDMA induces a transient period of heightened structural and functional plasticity in prefrontal circuits that coincides with fear extinction learning, providing mechanistic insight into its therapeutic potential for trauma-related disorders.

## Results

We first examined the impact of a single dose of MDMA on structural neuroplasticity in the cingulate/premotor (Cg1/M2) area of the mPFC (Fig 1A), a region in which the classical psychedelics psilocybin and 5-MeO-DMT have previously been shown to induce long-lasting increases in dendritic spine density for over 30 days (Shao et al., 2021; Jefferson et al., 2023). Using *in vivo* longitudinal two-photon microscopy of *Thy1^GFP^* mice, in which layer 5 and 6 pyramidal neurons are sparsely labeled with GFP, we imaged individual dendrites and dendritic spines at regular 2-day intervals starting 3 days prior to injection of 7.8 mg/kg MDMA and continuing until 7 days post-drug, with one additional session at the extended timepoint of 34 days post-drug (Fig 1B). Individual dendritic spines were tracked across sessions and categorized as stable, new, or eliminated. We imaged 1,264 spines across 75 dendritic segments from 10 animals (6 male, 4 female). We found that a single dose of MDMA robustly increases spine density (MDMA: +19± 3.8% compared to saline: -4.9±2.4 on Day 1, mean ± SEM, Fig 1C) and spine density remains elevated up to at least 7 days post-injection (MDMA: +13±4.5% compared to saline: -5.8±2.4% on Day 7; main treatment effect, p = 0.044, Two-way ANOVA). Spine density returned to saline levels 34 days after injection (MDMA: +4.7±3.9% compared to saline: +1.3±3.1% on Day 34). As an increase in spine density could be explained by an elevated rate of new spine formation or decreased rate of spine elimination, we calculated formation and elimination rates for each dendritic segment by counting the number of new spines formed or existing spines eliminated relative to total spine number on the first imaging session. We found that MDMA increases the rate of new spine formation only on Day +1 (Fig 1D, p = 0.0004 Bonferroni’s test). In contrast, elimination rate is not significantly impacted by MDMA (Fig 1E, p>0.05, supplemental Fig 1), suggesting that increases in spine density are driven by a transient elevation of spine formation rate in the first day following drug delivery.

**Figure 1.**
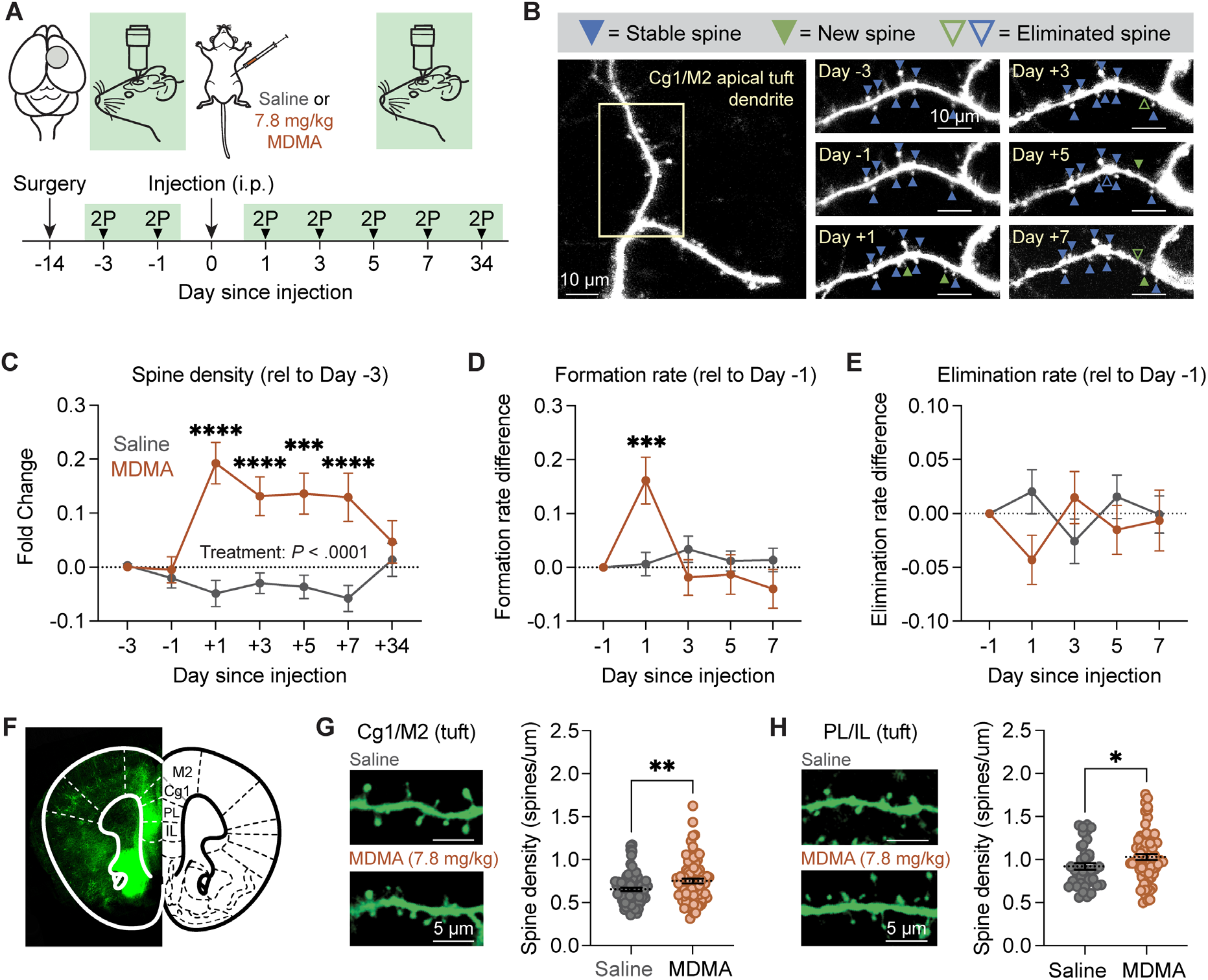
- MDMA enhances structural neuroplasticity in the medial frontal cortex. (A) Schematic and timeline of longitudinal two-photon imaging. (B) Example field of view (FOV) tracking dendritic spines across days in mouse treated with 7.8 mg/kg MDMA on Day 0. Scale bars are 10 µm. (C) Density of dendritic spines on apical tuft dendrites of layer V pyramidal neurons after MDMA (orange) or saline (gray), expressed as fold change from baseline on Day -3. Data shown as mean ± SEM. Two-way ANOVA: main effect of drug, F(1,469) = 58.12, p < 0.0001; effect of time, F(6,469) = 2.309, p = 0.033; drug x time interaction, F(6,469) = 6.101, p < 0.0001. (D) Spine formation rate expressed as a difference from baseline on Day -1 relative to Day -3. Two-way ANOVA: main effect of drug, F(1,341) = 0.07292, p = 0.7873; effect of time, F(4,341) = 4.001, p = 0.0035; drug x time interaction, F(4,341) = 5.009, p = 0.0006. Bonferroni’s multiple comparisons test: p = 0.0004 on Day 1, saline vs. MDMA. (E) Spine elimination rate expressed as a difference from baseline on Day -1 relative to Day -3. Two-way ANOVA: p > 0.05 for drug, time and drug x time interaction. (F) Example coronal section of mPFC from confocal imaging of *Thy1^GFP^*mouse. (G-H) Example FOVs of apical tuft dendrites from Cg1/M2 (G) and PL/IL (H) 24 hours after saline or MDMA and quantification of spine density, scale bars 5 μm. Two-tailed *t*-test: p = 0.0051 (G), p = 0.0449 (H).

Given recent interest in MDMA-assisted therapy for the treatment of PTSD, we wanted to determine whether MDMA also induces structural neuroplasticity in frontal cortical regions involved in the extinction of learned fear. To this end, we performed confocal imaging of dendritic spines in multiple mPFC subregions including Cg1/M2, prelimbic and infralimbic (PL/IL) cortices of *Thy1^GFP^* mice 24 hours after injection of MDMA (Fig 1F-H). We first confirmed that MDMA increases spine density in tuft dendrites in Cg1/M2 (Fig 1G, p = 0.0051, *t*-test), as well as basal dendrites in this region (supplemental Fig 2, p = 0.04, t-test), consistent with our two-photon imaging data. We found that MDMA increases spine density in both tuft (Fig 1H, p = 0.045, t-test) and basal (supplemental Fig 2, p = 0.031, t-test) dendrites in the PL/IL. Overall, we found that a single injection of MDMA increased structural neuroplasticity by promoting the formation of new dendritic spines, similar to classic psychedelics, though with a shorter-lived effect (Shao et al., 2021; Jefferson et al., 2023). We additionally found that MDMA’s effects on dendritic spine density could be observed across mPFC regions involved in the acquisition and extinction of learned fear, suggesting that the increased structural neuroplasticity in fear-related regions could underlie the fear extinction enhancement effects of MDMA.

To further elucidate the effects of MDMA on synaptic structure, we performed quantitative proteomic analysis of mPFC synaptosomal fractions from mice treated with MDMA (7.8 mg/kg) 24 hours prior to tissue collection (Fig 2A), the same timepoint at which we observed increased dendritic spines in this region. We identified 206 proteins that were differentially expressed between MDMA and saline synaptosome fractions with an adjusted p-value <0.05 (Fig 2B). Of the differentially expressed proteins (DEPs) meeting this significance threshold, 126 were downregulated and 80 were upregulated. Gene Ontology (GO) analysis was used to identify biological processes enriched in the list of DEPs. We found that the top process regulated by MDMA was neurofilament bundle assembly (Fig 2C). Other processes impacted by MDMA included metabolism of serotonin and dopamine, positive regulation of synaptic transmission, and cation transmembrane transport. As neurofilament proteins have been shown to regulate synaptic transmission and receptor trafficking (Sainio et al., 2022; Kim et al., 2002), and levels within dendritic spines scale with synaptic strength (Gürth et al., 2023), we chose to validate changes in NF-heavy (NF-H), NF-medium (NF-M), and NF-light (NF-L) subunits in synaptosome fractions by western blot (Fig 2D). We found that levels of NF-H, NF-M and NF-L were all significantly upregulated in MDMA-treated mice relative to saline-treated mice (p = 0.029, 0.042, and 0.048, respectively, t-test), consistent with a sustained effect of the drug on synaptic structure at the molecular level. Ingenuity Pathway Analysis was also performed to identify canonical pathways that were significantly enriched in this list of MDMA-induced DEPs (Fig 2E). The most significantly altered pathways encompassed multiple processes related to translation including initiation, elongation, and termination, suggesting that local translation at the synapse is upregulated 24 hours after MDMA. ROBO receptor signaling, which is known to promote synaptogenesis, was also upregulated after MDMA. Overall, these data are consistent with plasticity on the subcellular level after MDMA administration.

**Figure 2.**
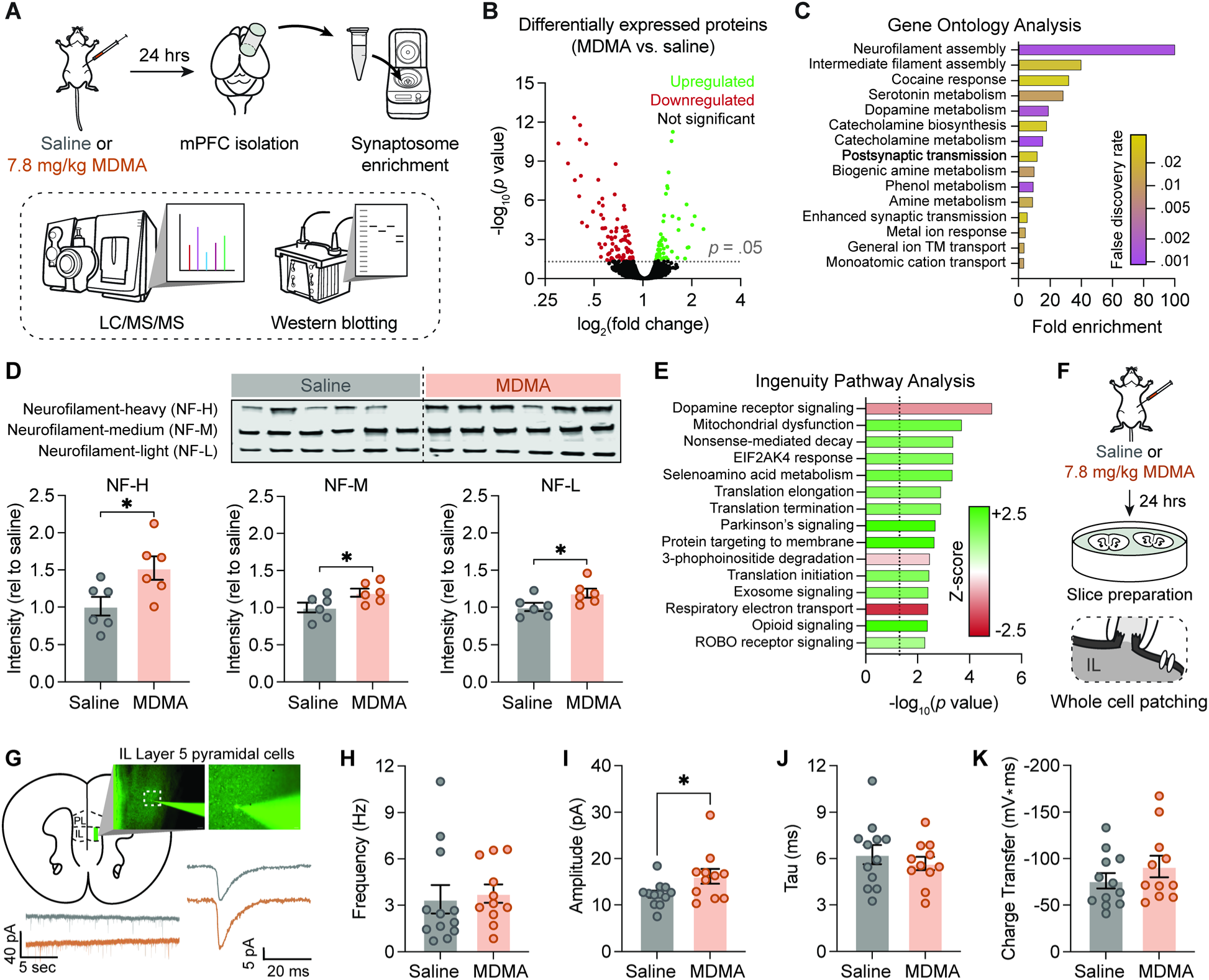
- MDMA alters the synaptic proteome and enhances excitatory postsynaptic strength in mPFC. (A) Schematic of synaptosome isolation from mPFC 24 hours after saline (gray) or MDMA 7.8 mg/kg (orange) followed by mass spectrometry and western blotting. (B) Volcano plot of differentially expressed proteins (DEPs) with proteins meeting significance threshold of p < 0.05 color-coded for upregulation (green) or downregulation (red) with MDMA. (C) Gene ontology analysis of DEPs following MDMA showing biological processes with highest fold enrichment, color-coded by false discovery rate. (D) Representative western blots and quantification for neurofilament-heavy (NF-H), neurofilament-medium (NF-M), and neurofilament-light (NF-L) proteins in synaptosomal fractions. Quantification expressed as relative to saline average. Two-tailed *t*-test: p = 0.029 (NF-H), p = 0.042 (NF-M), p = 0.048 (NF-L). (E) Ingenuity Pathway Analysis (IPA) for top pathways regulated by MDMA sorted by -log_10_(p-value) and color-coded by Z-score (green = predicted upregulation, red = predicted downregulation). (F) Schematic summarizing the preparation of acute brain slices containing the mPFC 24 hours after i.p. injection of saline or MDMA (7.8 mg/kg). (G) (Top) Representative image of a patched neuron imaged with DIC (Olympus BX51WI) in low magnification (10X/0.25 NA) (Left) and higher magnification of boxed region (40X/0.80 NA) (Right). Scale bars: left, 200 µm; right, 40 µm. (Bottom) Example voltage clamp traces from saline (grey) or MDMA (orange) animals (bottom left) and averaged sEPSC traces (bottom right). (H-K) Histograms comparing sEPSC frequency, amplitude, tau, and charge transfer. *p < 0.05, unpaired Student’s t-tests. N = 23 neurons from 8 mice (4 male, 4 female).

To investigate the functional outcomes of MDMA-induced spinogenesis and neuronal plasticity, we recorded IL cortex layer 5 (L5) pyramidal neurons using whole cell patch clamp electrophysiology 24 hours after injection of either saline or MDMA (Fig 2F, 7.8 mg/kg, IP). Cells were identified based on morphology and membrane properties. When measuring glutamatergic synaptic transmission, we found that MDMA-treated mice had significantly larger spontaneous excitatory postsynaptic current (sEPSC) amplitudes when compared to saline controls (t_21_ = 2.235, p = 0.0364, unpaired Student’s t-test, Fig 2I). sEPSC frequency (t_21_ = 0.3267, p = 0.7471, unpaired Student’s t-test, Fig 2H), tau (t_21_ = 0.7463, p = 0.4637, unpaired Student’s t-test, Fig 2J), and charge (t_21_ = 1.102, p = 0.2830, unpaired Student’s t test, Fig 2K) were unchanged in MDMA animals. Membrane properties such as resting membrane potential (RMP) and input resistance (R_in_) were also not significantly different between MDMA and saline animals (data not shown). These results suggest increased postsynaptic sensitivity to glutamate after MDMA experience rather than presynaptic modulation, supporting findings of structural dendritic plasticity after MDMA.

Decades of research have shown a link between memory encoding, recall, and synaptic connectivity (Bailey et al., 1989; Bailey & Kandel, 1993; Yang et al., 2009; Sanders et al., 2012). Thus, we next investigated how MDMA-induced neuroplasticity would affect the neuronal representation of conditioned auditory fear responses over several days. First, we examined how MDMA changes fear extinction during a prolonged, 5-day protocol (Fig 3A), based on previous studies showing enhanced fear extinction after MDMA administration (Young et al., 2015). On day 1 (conditioning), we presented mice (n=24) with a single 30-second auditory cue that co-terminated with a 1-second foot shock. 24 hours later, mice were randomly assigned to either MDMA (7.8 mg/kg) or saline (n=12 per group). 30 minutes after drug injection the mice were placed back in the behavior recording apparatus with different contextual cues and presented with 6 cues without shock for fear extinction training. We continued the extinction testing, with 8 cues on day 3, and 10 cues on days 4 and 5. Consistent with previous findings (Young et al., 2015; 2017), mice treated with MDMA before the extinction training session exhibited reduced freezing behavior in subsequent extinction testing days (Fig 3B-C).

**Figure 3.**
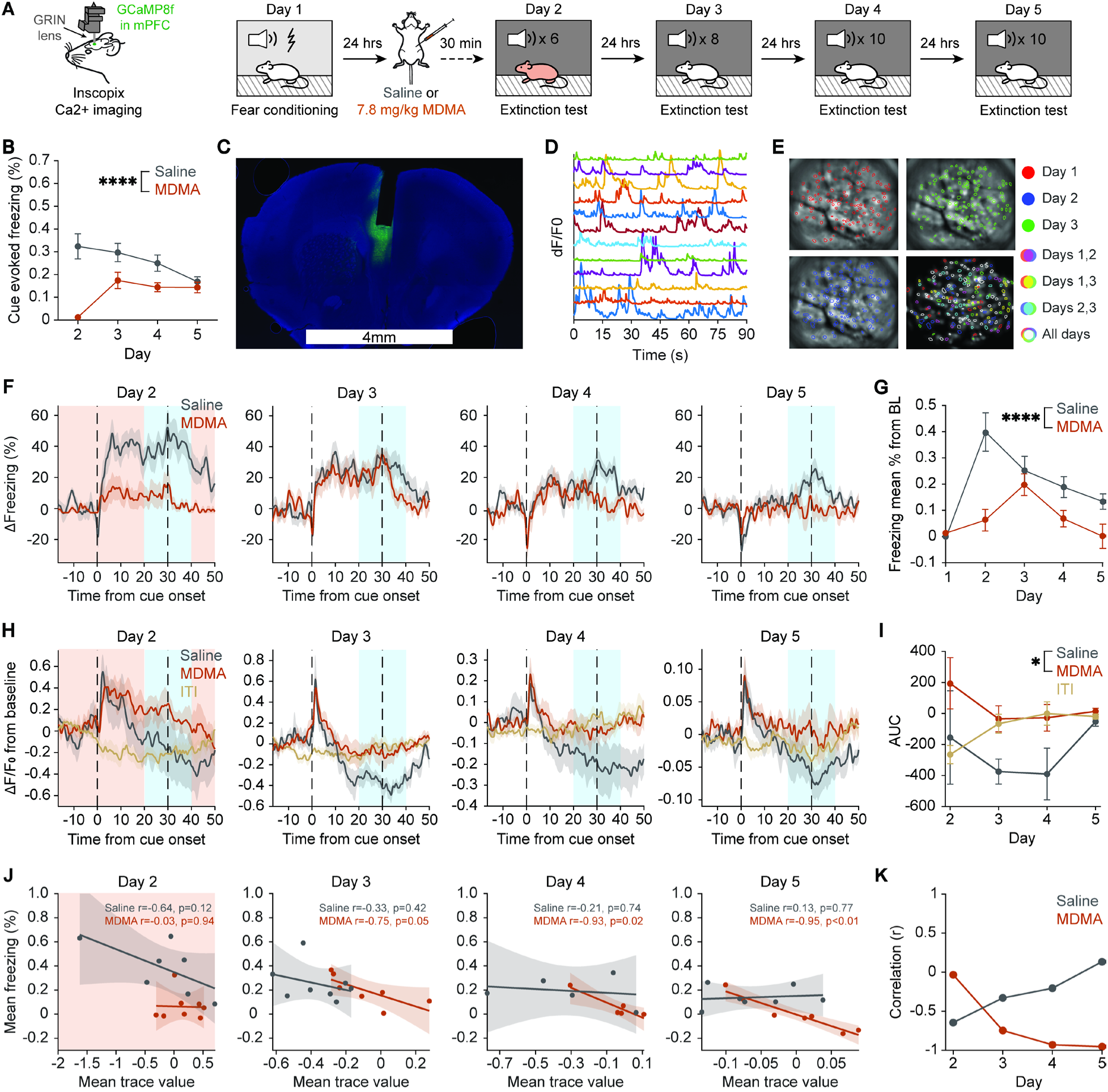
- MDMA alters population Ca²⁺ responses to a learned auditory cue during extinction. (A) Schematic of the fear extinction protocol performed during infralimbic (IL) Ca²⁺ imaging. Mice underwent daily extinction sessions consisting of repeated presentations of the conditioned auditory cue while Ca²⁺ activity was recorded. (B) MDMA-treated mice exhibit reduced freezing during extinction. Mean freezing during the 30-s auditory cue across extinction days (n=12 mice per group; 2-way ANOVA: effect of Treatment, F(1,88) = 37.23, p<0.0001; effect of Time, F(3,88) = 2.330, p = 0.08; Time × Treatment interaction, F(3,88) = 6.828, p=0.0003). (C) Representative histology showing lens placement in IL, scale bar = 4mm. (D) Example deconvolved Ca2+ traces from 12 cells identified in a single session. (E) Example of longitudinal cell registration demonstrated across three imaging days. Cellular footprints extracted for each day are shown overlaid on raw fluorescence images, with matched cells color-coded across sessions using IDPS-based registration. (F) Freezing values plotted for each timepoint after baseline normalization by subtracting each mouse’s mean freezing during the 30 s preceding cue onset. Data are plotted as mean ± SEM (n=8 mice). (G) Same as in (B), but for freezing measured from 10 s before to 10 s after cue offset in the implanted cohort (2-way ANOVA: effect of Treatment, F(1,56) = 23.87, p<0.0001; effect of Time, F(3,56) = 5.79, p = 0.0016; Time × Treatment interaction, F(3,56) = 3.455, p=0.022). (H) Baseline normalized population Ca2+ responses (mean ΔF/F across active neurons for each day) reveal a biphasic response in saline-treated mice across extinction testing days, consisting of an early cue onset excitation followed by a late inhibition. (I) Late cue responses (AUC of the mean trace from 10 s before to 10 s after cue offset) are inhibited in saline, but not in MDMA-treated mice (mixed effects model: effect of Treatment, *F*(1,48)= 6.30, *P* = 0.015; no main effect of Time, *F*(1.43,22.84)=1.21, *P*=0.30, or Time × Treatment interaction, *F*(1.43,22.84)=0.44, *P*=0.59). Data are plotted as mean ± SEM. (J-K) Decreased mPFC Ca^2+^ activity around cue offset is correlated with increased mouse freezing. (J) Correlation between the mean value of the population Ca2+ response (H) and mean baseline reduced freezing (F) during the 20 second window around cue offset shows a strong negative correlation in only in MDMA-treated mice in extinction testing days (3-5). Each data point represents the mean values for each mouse. Shaded areas are 95% confidence intervals for fitted regression. (K) Correlation coefficients across days (from J) show that freezing and GCaMP signal become decorrelated over extinction days in saline-treated mice, but remained strongly negatively correlated in MDMA-treated mice.

Next, we utilized longitudinal miniscope Ca^2+^ imaging to record the activity of neuronal ensembles in the IL of another cohort of mice that underwent the same fear extinction protocol. We recorded activity from individual cells (Fig 3D-E, n=38-130 cells per session) and registered the cells across days of extinction training and testing (n=78-246 cells per mouse) while simultaneously assessing freezing behavior. We then analyzed freezing behavior at high temporal resolution to determine how responses to the learned auditory cue diverged between treatment groups. We found that much of the treatment-level difference in behavior occurred around the end of the 30 second cue (Fig 3 F-G), and that this difference increased across extinction learning. We then calculated the mean Ca^2+^ activity trace across all recorded cells for each mouse and revealed that both MDMA- and saline-treated mice showed an immediate increase in activity at cue onset across all extinction days (Fig 3H). Notably, the cue-evoked Ca^2+^ trace in saline-treated mice showed a biphasic response; both MDMA- and saline-treated mice showed an immediate increase in activity at cue onset, but only saline-treated mice showed a reduction in Ca^2+^ activity later in the cue. MDMA-treated mice did not exhibit this late-cue negative deflection; instead, activity throughout the late cue period returned to pre-cue baseline (Fig 3H-I).

We then asked whether the late-cue negative deflection seen in the neuronal activity traces correlates to the freezing behavior. For each extinction day, we calculated the mean value of the freezing and GCaMP traces for each mouse across all cue presentations (Fig 3J). Consistent with the known role of IL in suppression of freezing behavior (Milad & Quirk, 2002; Do-Monte et al., 2015), GCaMP activity in the saline-treated group negatively correlated to freezing during extinction training (day 2). This relationship disappeared on subsequent days of extinction testing (days 3-5). The MDMA-treated group, however, maintained this negative correlation throughout the extinction testing days (Fig 3J-K, supplemental Fig 4).

Next, we investigated how MDMA-induced synaptic changes altered individual cells’ responses to the learned auditory cue. Using the IDPS cell registration algorithm, we registered the recorded cells across all days (Fig 3E, n=78-246 cells per mouse). The registered number of cells per mouse, the mean registration score per cell, and mean centroid distances across days were similar for the two treatment groups (p= 0.5, 0.43, and 0.88, respectively, supplemental Fig 5). Individual neuronal tuning curve responses varied across cue presentations (Fig 4A). Thus, we measured the similarity of IL tuning curves across cue presentations by computing pairwise Pearson correlations of Ca^2+^ traces of individual cells directly after the cue onset (within the first 7 seconds) across different cues over all extinction and testing (Fig 4B). We examined how MDMA-induced plasticity affected long-term differences in IL activity by calculating the similarity of the ensemble responses as a function of days of separation between all cue-pairs (Rubin et al., 2015; Geva et al., 2023). We found that the similarity between cue responses across different days was significantly lower in MDMA-treated mice than in the saline-treated group (Fig 4C).

**Figure 4.**
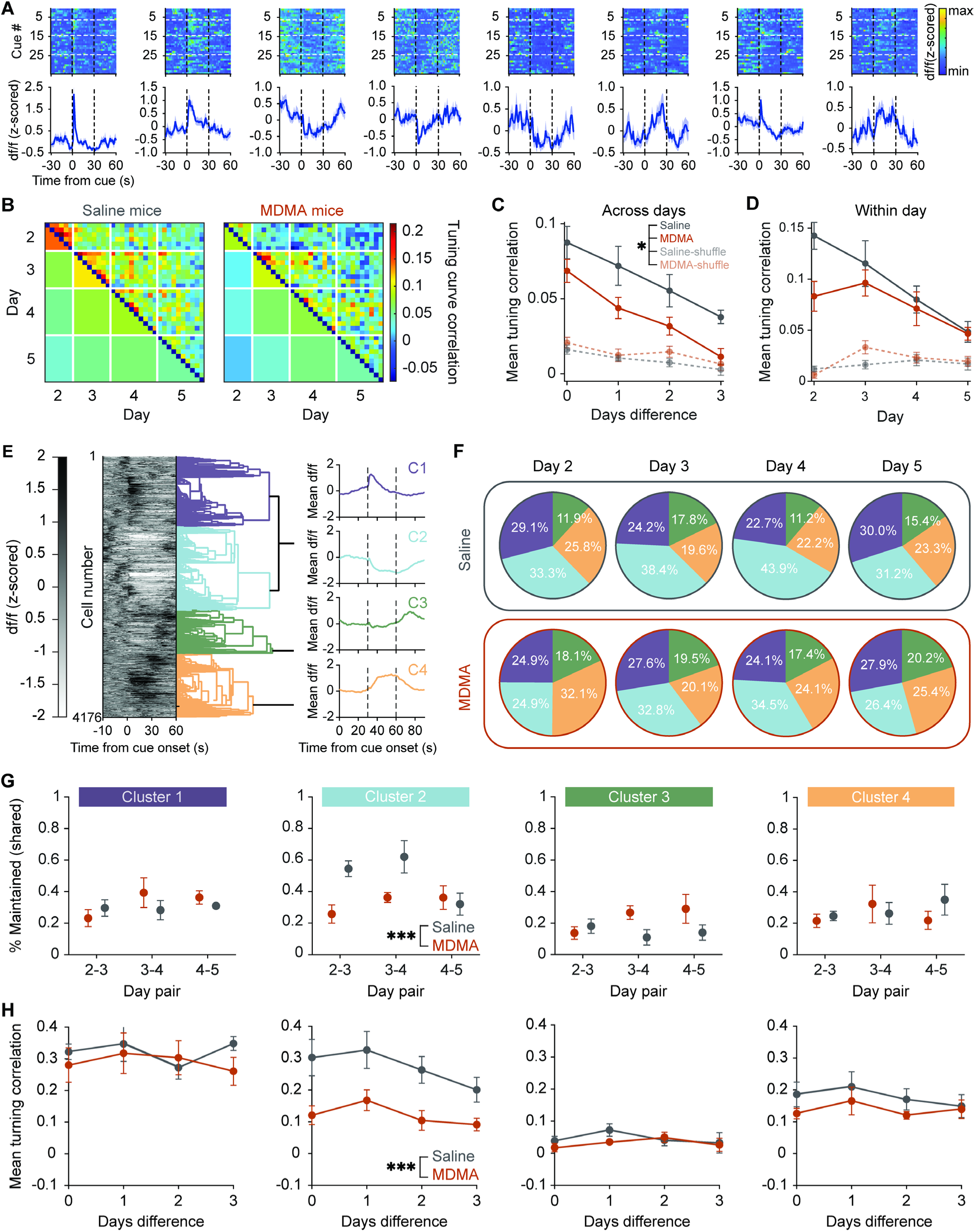
- MDMA enhanced plasticity increases representational drift in IL during fear extinction. (A) Examples of individual cell responses to all cue presentations throughout extinction training, plotted for each cell from 30 seconds before cue onset to 30 seconds after cue offset, top – heatmap of GCaMP traces for each cue presentation, bottom – mean ±SEM across all cues. (B) Mean tuning curve correlations of individual cells’ cue evoked Ca^2+^ traces (within 7s of cue onset) across cue-pairs for Saline (left) and MDMA (right) treated mice. Areas under the main diagonal show mean correlation across cues for corresponding day pairs. (C) Pairwise correlations of tuning curves of individual cells (within 7s of cue onset) as a function of time difference between cue presentations, averaged per mouse. MDMA-treated mice show lower ensemble tuning curve correlations compared to saline-treated mice across all time differences, indicating faster representational drift (Mixed-effects model: main effect of treatment, F(1,14) = 6.322, p = 0.0248; effect of time, F (2.012, 25.49) = 23.38, p<0.0001; time × treatment interaction, F (2.012, 25.49) = 0.1783, p = 0.839). Dashed lines represent shuffle controls where cell identities were randomly permuted. Data shown as mean ± SEM. (D) Within day. Pairwise correlations of cue-evoked activity patterns for cue presentations within the same extinction test day (Days 2-5). MDMA-treated mice and saline-treated mice show similar within-day ensemble stability (Mixed-effects model: main effect of treatment, F (1, 14) = 2.455, p = 0.14; effect of day, F (1.736, 19.68) = 10.72, p=0.001; day × treatment interaction, F (1.736, 19.68) = 2.040, p = 0.16). Dashed lines represent shuffle controls where cell identities were randomly permuted. Data shown as mean ± SEM. (E) Dendrogram showing clustering of cellular responses across all days. For each day responses were averaged across all cue presentations for each cell (left). Mean tuning curve for the 4 clusters (right). (F) Proportions of cells in each cluster on each day for saline (top) and MDMA (bottom) treated mice. (G) Percentage of cells that maintained the same cluster identity on consecutive day pairs for each response cluster. A linear mixed-effects model revealed a significant main effect of treatment (*F*(1,29)=15.27, p=0.0005) and a day pair × treatment interaction (F(2,29) = 6.49, p = 0.005) for cluster 2 only, indicating that MDMA-treated mice had lower cluster stability in this subpopulation. No significant effects of treatment were observed for clusters 1, 3, or 4 (p > 0.05 for all terms). (H) Pairwise correlations of tuning curves of individual cells (over 30 seconds of cue presentation) for cells identified in different clusters, as a function of time difference between cue presentations, averaged per mouse. A linear mixed-effects model revealed a significant main effect of treatment (F(1,56) = 11.30, p = 0.001), and time difference (F(1,56) = 8.21, p = 0.006) for cluster 2 only, with MDMA-treated mice showing lower tuning curve correlations, indicating faster representational drift for cluster 2 neurons. No significant effects of treatment or time lag were observed for clusters 1, 3, or 4 (p > 0.05 for all terms). Data shown as mean ± SEM.

While these ensemble-level analyses reveal that MDMA accelerates representational drift overall, they do not distinguish whether this effect is uniform across all cells or driven by specific subpopulations. To address this, we used hierarchical clustering to identify distinct response patterns and asked whether MDMA differentially affected the stability of each cluster (Fig 4E-F, see methods). The four response patterns we found correspond to a cue-onset excitation (cluster 1), cue-onset suppression (cluster 2), post-cue offset response (cluster 3), and a sustained excitation of Ca^2+^ activity after cue onset (cluster 4). We then asked whether each neuron maintains its response pattern across days and found that only cluster 2 significantly differed in day-to-day stability between treatment groups, with more cells maintaining cluster identity across days 2-to-3 and 3-to-4 (Fig 4G) in the saline-treated mice. Finally, we computed the tuning curve correlations (similarly to Fig 4B-D) for the entire duration of the 30-second auditory cue for cells that maintained their cluster identity across day pairs (Fig 4H). Again, only cluster 2 significantly differed in response similarity between treatment groups. Taken together, these results demonstrate that MDMA induces a faster rate of representational drift in cue evoked responses in IL, suggesting a relation between this phenomenon and the observed changes in fear-related behavioral responses.

## Discussion

Our findings demonstrate that a single dose of MDMA robustly increases dendritic spine density in the mouse mPFC (Fig 1), adding MDMA to the growing list of rapid acting psychoactive compounds, including psilocybin, 5-MeO-DMT, and ketamine, that induce structural neuroplasticity in frontal cortical circuits (Shao et al., 2021; Jefferson et al., 2023; Moda-Sava et al., 2019; Muñoz-Cuevas et al., 2013). However, we find a key difference in the temporal dynamics of these structural changes. While psilocybin and 5-MeO-DMT produce spine density increases that persist for over 30 days (Shao et al., 2021; Jefferson et al., 2023; Shao et al., 2025), MDMA-induced spinogenesis is more transient, with spine density remaining elevated for at least 7 days but returning to baseline by 34 days after injection (Fig 1C). While repeated cocaine exposure has been shown to enhance spine turnover in the frontal cortex (Muñoz-Cuevas et al., 2013), the effects of MDMA induced spinogenesis are distinct from both cocaine and classical psychedelics. The difference in the duration of structural neuroplasticity is notable given the distinct pharmacological profiles of these drugs. Classic serotonergic psychedelics act primarily as direct agonists of the 5-HT2A receptor, whereas MDMA mainly acts as a monoamine releasing agent (Torres et al., 2003). Recent work has shown that MDMA exhibits weak partial agonism at 5-HT2A receptors *in vitro*, with changes in dendritic spine density being at least partially dependent on the 5-HT2A receptor (Gaines-Smith et al., 2026). These findings suggest that the effects of MDMA and serotonergic psychedelics on structural plasticity may converge on the 5-HT2A receptor, but the magnitude and duration of receptor engagement could differ substantially, leading to differences in spine dynamics over time. Future work employing selective 5-HT2A antagonists or receptor knockout models during drug administration will be necessary to definitively parse the contributions of direct partial agonism, indirect receptor activation via released serotonin, and 5-HT2A-independent mechanisms through other 5-HT receptors, dopamine or norepinephrine signaling.

Importantly, MDMA-induced spinogenesis was not limited to specific subregions of the mPFC, with increased spine density observed in Cg1/M2, PL/IL cortices, in both tuft and basal dendrites. This lack of regional selectivity suggests that MDMA does not by itself bias the circuit toward extinction but rather opens a window of structural plasticity that may allow experience-dependent processes, such as extinction training, to alter connectivity in behaviorally relevant circuits. This interpretation aligns with the clinical model of MDMA-assisted therapy, in which the drug is not administered in isolation but in the context of guided psychotherapy, and is consistent with prior work demonstrating that MDMA enables plasticity of social reward learning (Nardou et al., 2023). Future studies might examine whether MDMA also enhances spinogenesis in other fear-related brain regions, such as the amygdala or hippocampus, and whether such structural plasticity affects circuit function.

Our findings demonstrate concurrent evidence that MDMA induces postsynaptic plasticity in mPFC pyramidal neurons using different methods and levels of analysis. Structurally, we observed increased spine density in both tuft and basal dendrites in multiple mPFC subregions (Fig 1F-H, supplemental Fig 2). At the molecular level, quantitative proteomic analysis of mPFC synaptosomal fractions revealed upregulation of proteins related to neurofilament bundle assembly, monoamine metabolism and synaptic transmission (Fig 2C-E). Neurofilament proteins are integral cytoskeletal components of synapses whose levels in the dendritic spine have been shown to scale with synaptic strength (Gurth et al., 2023). Deletion of these proteins has been demonstrated to significantly impair hippocampal long-term potentiation (LTP) (Yuan et al., 2015), suggesting that they are essential for induction of functional plasticity in that region. Regarding functional synaptic plasticity, we found that MDMA increases the amplitude of sEPSCs in IL L5 pyramidal neurons, without altering their frequency, decay kinetics, or charge (Fig 2G-K). The selective increase in sEPSC amplitude, in the absence of frequency changes, is consistent with a postsynaptic mechanism, rather than alterations to presynaptic release probability. Though no prior studies have observed the effects of MDMA on mPFC whole-cell slice electrophysiology, our findings support literature that MDMA can facilitate plasticity in the brain. Acute application of MDMA enhances LTP in rat CA3-CA1 hippocampal field excitatory post synaptic potentials (Mlinar et al., 2008, Rozas et al., 2012; Mlinar et al., 2015). In the nucleus accumbens, acute MDMA application has been shown to induce long-term depression (LTD) of electrically evoked EPSCs in both D1 and D2 medium spiny neurons (Heifets et al., 2019). Rats that self-administered MDMA on a fixed ratio schedule showed increased central amygdala neuron miniature inhibitory postsynaptic current (mIPSC) amplitude the day after the final self-administration session (Khom et al., 2021), suggesting MDMA-induced postsynaptic long-term plasticity in GABAergic circuits. Thus, our results from electrophysiology and proteomics experiments reinforce the hypothesis of a postsynaptic locus for the strengthening in mPFC synaptic connections.

A central question we addressed in our study is how structural plasticity in the mPFC translates to altered circuit function during fear extinction. Fear extinction is widely used as a preclinical model for exposure-based therapies used to treat PTSD (Milad & Quirk, 2012, Paredes et al., 2019), making IL circuit dynamics during extinction directly relevant to understanding MDMA’s therapeutic potential. Understanding how MDMA-induced plasticity alters IL circuit dynamics during extinction may therefore provide mechanistic insight into how the drug facilitates therapeutic outcomes. We found that MDMA strengthened the coupling between IL ensemble activity and freezing suppression across extinction testing days, a pattern consistent with enhanced top-down inhibitory control over fear expression. Interestingly, saline-treated mice showed a late-cue inhibitory deflection in IL activity that correlated with freezing suppression on day 2 but decoupled over subsequent extinction days, suggesting that this response pattern may reflect an early extinction signal that is not maintained without the additional support provided by MDMA-induced spinogenesis. While the temporal cooccurrence of structural and functional changes is suggestive, the causal relationship between spinogenesis and altered ensemble dynamics remains correlational. Future experiments selectively blocking spine formation during MDMA administration would be necessary to establish whether spinogenesis is required for the observed functional changes.

Our finding that MDMA-treated mice show lower ensemble tuning curve correlations across days demonstrates that MDMA-induced spinogenesis is associated with accelerated representational drift in IL during fear extinction. Representational drift, the gradual evolution of neural population codes over time, has been observed in multiple cortical regions and is thought to reflect ongoing synaptic reorganization (Rubin et al., 2015; Geva et al., 2023; Rule et al., 2019; Driscoll et al., 2022; Deitch et al., 2021). In our context, the increased drift in MDMA-treated mice suggests that the IL ensemble encoding of the conditioned cue is more rapidly updated across days. Notably, at the population level, MDMA strengthened the coupling between IL ensemble activity and freezing suppression across extinction days (Fig 3J-K). At the single-cell level, the accelerated drift was strongest in a functionally defined subpopulation showing inhibitory responses to the fear cue (Fig 4G-H), suggesting that MDMA-induced plasticity preferentially reorganizes the representations of neurons most directly engaged in signaling threat suppression. This observation supports models that suggest that representational drift is at least partially driven by gradual changes in synaptic connections (Kalle Kossio et al., 2021; Devalle et al., 2025). Additionally, our results support the interpretation of representational drift as a signature of continued learning, rather than mere noise or degradation, as MDMA-treated mice show progressively reduced freezing across the same time period over which their ensemble representations are evolving most rapidly. These findings also highlight the utility of pharmacological neuroplasticity as an experimental tool for probing the relationship between synaptic reorganization and representational drift, offering a tractable approach for future studies seeking to causally link structural and functional plasticity in cortical circuits.

An intriguing aspect of our data concerns the temporal interaction between MDMA’s structural and functional effects. Structurally, the peak of new spine formation occurs within 1 day, yet behaviorally, the divergence in freezing between MDMA and saline groups becomes increasingly pronounced over the subsequent days of extinction training. Similarly, our analysis of representational similarity in IL reveals that the divergence in ensemble dynamics between treatment groups is most apparent across days rather than within a single session. This temporal dissociation suggests that the initial wave of spinogenesis sets up the conditions for ongoing learning, rather than directly encoding extinction memories at the time of drug administration. Thus, we suggest that the structural changes induced by MDMA within the first 24 hours serve as the substrate upon which continued extinction learning is inscribed over the following days, consistent with a model in which the drug facilitates learning not by creating new memories itself but by enhancing the brain’s capacity for synaptic reorganization in response to experience.

## Conclusions

Taken together, our results reveal that MDMA induces a period of enhanced structural and functional plasticity in the mPFC that facilitates fear extinction learning. The convergence of structural, molecular, electrophysiological, and functional imaging data provides a multi-level mechanistic framework for understanding how MDMA may facilitate the updating of fear memories. We propose that MDMA, through the formation of new dendritic spines, opens a plasticity window which can then enable experience-dependent processes to reshape cortical representations of learned threats.

## Methods

### Animals

All experiments were approved by Yale University’s Institutional Animal Care and Use Committee (IACUC).

For spine counting experiments we used lab-bred heterogenous Thy1^GFP^ line M mice (Tg(Thy1-EGFP)MJrs/J, Stock #007788, Jackson Laboratory) crossed with C57BL/6J (Stock No. 000664, Jackson Laboratory). For Two-photon imaging experiments we used mice aged 8-12 weeks at time of surgeries. For confocal imaging experiment we used 17-32 weeks old mice. For synaptosome proteomics,

For electrophysiology and miniscope imaging experiments we used C57BL/6J mice bought directly from Jackson laboratories, aged 8-12 weeks at time of first surgeries. All animals were group housed (2-5 mice per cage), kept on a 12h/12h light-dark cycle, and provided ad libitum chow and water. All experiments were performed during the light cycle (7:00-19:00).

### Surgeries

For all surgeries we first injected all mice with carprofen (5 mg/kg, s.c.; 024751, Henry Schein Animal Health) and dexamethasone (3 mg/kg, i.m.; 002459, Henry Schein Animal Health) immediately prior to surgery. We then anesthetized the mice using isofluorane (∼3% for induction, ∼1% for maintenance) and head-fixed them to a stereotaxic apparatus with a heating pad set to 38 °C under the body, and lubricated their eyes with ophthalmic ointment. We then clipped scalp hair and cleaned the skin with betadine and ethanol before removing the scalp and connective tissue and cleaning the skull using hydrogen peroxide.

For two-photon imaging we created a ∼3 mm circular craniotomy above the right medial frontal cortex (center position: + 1.5 mm anterior-posterior, AP; + 0.4 mm medial-lateral, ML; relative to bregma) using a dental drill. We then placed a glass window, consisting of two round 3mm-diameter #1 thickness glass coverslips (64–0720 (CS-3R), Warner Instruments), bonded by UV-curing optical adhesive (NOA 61, Norland Products), over the craniotomy and secured it to the skull using adhesive (Henkel Loctite 454). We then affixed a stainless steel headplate to the skull using C&B Metabond (Parkell).

For miniscope imaging experiments, we performed a craniotomy (∼0.5-0.7 mm in diameter) with a dental drill above the mPFC at coordinates 1.65 mm anterior, 0.3 mm lateral, and 2.25 mm ventral to bregma. We injected 500 nL of AAV1.Syn.GCaMP8f virus (Addgene, 1.9 x 10¹³ particles per mL) using an automated microinjector (Nanoject, Drummond Scientific). We then lowered a 0.5 x 4 mm GRIN lens (ProView, Inscopix) to just above the injection site and secured it to the skull using super-glue (Loctite). We sealed the exposed areas of the skull with Metabond (Parkell).

After surgery, we returned the mice to their home cages for recovery. We monitored recovery over three days and administered carprofen (5 mg/kg, i.p.) and dexamethasone (3 mg/kg, i.p.) daily for 2-3 days following surgery. Mice recovered for at least 10 days after the surgery prior to the start of imaging experiments.

### Two-photon imaging

The excitation laser source was a Ti:Sapphire ultrafast femtosecond laser (Axon, Coherent), with intensity controlled by a Pockels cell (350-80-LA-02, Conoptics) and a shutter (LS6ZM2, Uniblitz via Vincent Associates). The beam was directed into a two-photon microscope (Movable Objective Microscope) that included a water-immersion high-numerical aperture objective (XLUMPLFLN, 20X/0.95 NA, Olympus). Excitation wavelength was 920 nm and emission from 475 to 550 nm was collected using a GaAsP photomultiplier tube (H7422-40MOD, Hamamatsu). The laser power measured at the objective was ≤ 40 mW. The two-photon microscope was controlled by ThorImageLS software (Thorlabs)

During imaging sessions, mice were head-fixed and lightly anesthetized with 1-1.5% isofluorane. Imaging sessions typically lasted ∼30 min and did not exceed 1h. Apical tuft dendrites were imaged at 0–400 μm below the dura. For Cg1/M2, we imaged within 0–400 μm of the midline as demarcated by the sagittal sinus. 1–10 fields of view were collected from each mouse, taking 10–40 μm-thick image stacks at 1 μm steps at 1024 × 1024 pixels with 0.642 μm per pixel resolution. The same imaging parameters were used for every imaging session. To image the same fields of view across multiple days, we would identify and return to a landmark on the top (12:00) of the glass window. Each mouse was imaged on days -3, -1, 1, 3, 5, 7, and 34 relative to the day of treatment. On the day of treatment (day 0) mice were injected with MDMA (7.8 mg/kg, i.p.) or saline.

### Two-photon imaging analysis

Images were analyzed using ImageJ (ref) with StackReg plug-in (ref) for motion correction. Protrusion of >0.4 μm from the dendritic shaft were considered dendritic spines. Change in spine density across days was calculated as a fold-change relative to the value measured on the first session (day -3) for that dendritic segment. Spine formation rate was calculated as the number of new spines formed between two consecutive imaging sessions divided by the total number of dendritic spines seen in the first imaging session. Spine elimination rate was calculated as the number of dendritic spines lost between two consecutive imaging sessions divided by the total number of dendritic spines seen in the first imaging session. The data were analyzed by an experimenter blind to drug condition.

### Confocal spine imaging

We randomly assigned mice from each cage were to receive MDMA (7.8 mg/kg, i.p, n = 2 males, 2 females) or saline (10 ml/kg, i.p., n = 2 females). 24 hours after injection, we deeply anesthetized the mice (isoflurane, 029405, Covetrus) and transcardially perfused them with 0.1M phosphate buffered saline (PBS, P4417, Sigma-Aldrich), followed by 4% paraformaldehyde (PFA, 15714, Electron Microscopy Sciences) in PBS, then collected and stored the brains in 4% PFA at 4°C, for 48 hours. We then sectioned the brains into 50-um thick slices using a vibratome (VT1000S, Leica) and mounted them onto coverslipped slides (71861-055, Electron Microscopy Sciences; 12-550-143, Fisherbrand).

We imaged the brain slices using a confocal microscope (SP8 Gated STED, Leica) 0-5 months after brain fixation. We acquired whole-slice images using a 20x air objective, and dendritic spines using a 63x oil objective, referenced to secondary motor cortex (M2) and prelimbic (PL)/infralimbic (IL) cortex using Paxinos and Franklin’s mouse brain atlas.

### Synaptosome isolation

Mice were euthanized and the mPFC including ACC, PL and IL were dissected on ice. Tissue was homogenized in TEVP-320 sucrose buffer (10 mM Tris-HCl, pH 7.4, 5 mM NaF, 1 mM Na_3_VO_4_, 1 mM EDTA, 1 mM EGTA, and 320 mM sucrose with EDTA-free cOmplete mini protease inhibitor mixture (11836170001; Roche Diagnostics)) and centrifuged at 1,000 x g for 10 min at 4 °C and supernatant collected (S1). S1 was centrifuged at 10,000 x g for 15 min at 4 °C to obtain a crude synaptosomal fraction (P2) and cytosolic fraction (S2). P2 fractions were resuspended and lysed in TEVP-35.6 sucrose buffer (10 mM Tris-HCl, pH 7.4, 5 mM NaF, 1 mM Na_3_VO_4_, 1 mM EDTA, 1 mM EGTA, and 35.6 mM sucrose with EDTA-free cOmplete mini protease inhibitor mixture (11836170001; Roche Diagnostics)) by pipetting and left on ice for 30 min. Protein concentration was determined using the *DC* protein assay (Bio-Rad).

### Mass spectrometry

Liquid chromatography and mass spectrometry was performed by the W. M. Keck Foundation Biotechnology Resource Laboratory, Yale School of Medicine, New Haven, CT. For proximity labeling experiments, only proteins that were enriched by FC ≥ 1.5 in samples compared to negative controls that were not injected with pAAV-BioID2-Shank3* were considered for analysis. For all experiments, only proteins present in >50% of samples and those identified by ≥ 2 unique peptides were included in differential expression analysis.

### Whole cell patch clamp electrophysiology

Mice were injected with MDMA (7.8 mg/kg, IP) or saline 24 hours prior to being sacrificed for whole cell patch slice recordings. Brains were rapidly removed, and coronal slices (300 μm-thick) containing the mPFC were cut using a vibratome (VT1200S; Leica Microsystems, Wetzlar, Germany) in an ice-cold aCSF cutting solution, containing the following (in mM): 93 NMDG, 2.5 KCl, 1.25 NaH_2_PO_4_, 30 NaHCO_3_, 20 HEPES, 25 glucose, 5 Na-ascorbate, 2 thiourea, 3 Na-pyruvate, 10 MgSO_4_, and 0.5 CaCl_2_, 300– 310 mOsm, pH 7.4 when continuously oxygenated with 95 % O_2_/5 % CO_2_. Slices were allowed to recover in the aCSF cutting solution at 34–36°C for 30 minutes during which increasing volumes of 2 M NaCl (up to a total of 1 mL NaCl/37.5 mL aCSF) were added every 5 minutes as previously described (Ting et al., 2018; Anderson et al., 2024). After recovery, the slices were transferred to a recording aCSF solution maintained at room temperature. Recording aCSF contained the following (in mM): 130 NaCl, 3 KCl, 1.25 NaH_2_PO_4_, 26 NaHCO_3_, 10 glucose, 1 MgCl_2_, and 2 CaCl_2_, pH 7.2–7.4, when saturated with 95 % O_2_/5 % CO_2_. For electrophysiology recordings, recording pipettes were pulled from borosilicate glass capillaries (World Precision Instruments, Sarasota, FL) to a resistance of 4–7 MΩ when filled with the intracellular solution. The intracellular solution contained the following (in mM): 145 potassium gluconate, 2 MgCl_2_, 2.5 KCl, 2.5 NaCl, 0.1 BAPTA, 10 HEPES, 2 Mg-ATP, and 0.5 GTP-Tris, pH 7.2–7.3 with KOH, osmolarity 280–290 mOsm. L5 IL neurons were viewed under an upright microscope (Olympus BX51WI) with infrared differential interference contrast optics and a 40x water-immersion objective. The recording chamber was continuously perfused (1–2 mL/min) with oxygenated recording aCSF warmed to 32 ± 1°C using an automatic temperature controller (Warner Instruments). L5 IL neurons were identified based on morphology. The patch pipette contained Alexa 488 and each patched neuron was imaged to verify location with a 10X objective after recording. To evaluate spontaneous excitatory postsynaptic currents (sEPSCs), neurons were voltage-clamped at −70 mV. To evaluate RMP, rheobase, and action potential firing, the cells were current-clamped and 500 ms depolarizing current steps were applied every second in 10 pA increments. All recordings were digitized at 20 kHz and lowpass-filtered at 2 kHz using a Digidata 1550B acquisition board (Molecular Devices, San Jose, CA) and pClamp11 software (Molecular Devices). Access resistance (10–30 MΩ) was monitored during recordings by injection of 10 mV hyperpolarizing pulses and data were discarded if access resistance changed >25 % over the course of data collection.

### Ca^2+^ imaging

At least 6 weeks after lens implantation we examined Ca^2+^ indicator expression and tissue health while mice were under light isoflurane anesthesia (0.8-1%). We selected for further imaging only those mice that exhibited healthy appearance of the tissue and clear cellular GCaMP activity. For the selected mice, we then attached the microscope baseplate using dental acrylic and light cured acrylic (Flow-It ALC, Pentron). We checked GCaMP signal further in awake animals before behavior experiment to ensure imaging quality. Only animals that showed clear cellular signals in at least 30 cells were used in the study. We then habituated the mice that were selected for the imaging experiment to human handling by allowing them to walk on the experimenters’ hands for at least two days prior to the start of experimental sessions.

### Fear conditioning

On Day 1, we placed the animals in a behavioral box (Med Associates - MED-VFC-OPTO-USB-M) with an added plastic opaque white wall to distinguish contexts. Additionally, we placed two drops of banana extract underneath the box. Together, this made context A. Before putting the mice in the behavior box, we recorded a 5-minute baseline of neuronal activity while the mice were in a clean cage near the behavior box. We began behavioral recordings after placing the mouse in the behavior box. 120 sec after the start of the recording, animals were presented with a 30-sec tone (79.5-80.5 dB, 5.5 kHz) culminating in a 1 second shock (1 mA). Recording was done at 30 Hz. The session ended 120 sec after the termination of the tone and shock pair. We cleaned the box with 70% ethanol between animals.

On Day 2, we first recorded a 5-minute baseline of neuronal activity. We then injected the mice with either 7.8 mg/kg MDMA or saline (i.p.) and recorded another 5 minutes of spontaneous activity outside the behavior box. We did not remove the miniature-microscope from the mouse throughout the session. 30 minutes after injection, we placed the mice in the same behavioral box without the plastic insert, and placed 2 drops of peppermint extract in the behavioral box to represent context B. We then recorded behavior and neuronal activity. 120 sec after the start of the recording, the mouse was presented with six 30-sec tones (79.5-80.5 dB, 5.5 kHz) with random inter-trial intervals of 45-75 sec. The session ended 120 sec after the termination of the sixth tone.

On Days 3-5, we again recorded 5 minutes of baseline activity before the animals were placed in the behavior box, set-up in context B. 120 sec after the start of the recording in the behavior box, the mouse was presented with eight tones on day 3, and 10 tones on days 4 and 5 (30-sec duration, 79.5-80.5 dB, 5.5 kHz) with random inter-trial intervals of 45-75 sec. The session ended 120 sec after the termination of the sixth tone.

Freezing behavior was measured using the VideoFreeze analysis program (Med Associates Inc.). Threshold for freezing behavior was manually set after each recording session to account for changes in output due to cable movement. Freezing behavior was extracted as a binary vector (0 non-freezing, 1-freezing) starting 30 seconds before each cue presentation and ending 30 seconds after cue offset. This vector was normalized by reducing the mean value of the 30 second pre-cue baseline from each timepoint of the 90 second trace. Behavior recording and calcium imaging were both done at 30 Hz, with a synchronization signal at the onset of each auditory cue.

### Calcium imaging processing

Cell activity and traces were extracted using routine IDPS (Inscopix) algorithms. After uploading, videos for each session were concatenated then spatially downsampled by a factor of 4 and spatially band-pass filtered. Videos were then motion corrected using high contrast areas of blood vessels as ROI references.

Cells were identified and extracted using CNMF-E (all parameters were set to the IDPS default, except the minimum pixel correlation, and minimum signal-to-noise ratio, which were set for each session separately at between 0.65-0.8, and 7-10, respectively). Extracted fluorescence traces from the CNMF-e were then deconvolved to extract estimated calcium events and traces. Cells across the 5 recording days were then longitudinally registered in IDPS (minimum normalized cross-correlation = 0.5). The registered deconvolved traces were then extracted for further analysis.

### Calcium activity trace analysis

Activity traces were parsed into segments of 90 seconds, beginning 30 seconds before the onset of each 30 second auditory cue and ending 30 seconds after cue offset. For each cell we then normalized the activity trace by reducing the value for each time point by the mean value of the trace during the 30 second pre-cue period. Mean activity traces were calculated as the mean value of the denoised baseline corrected while ignoring cells that were not identified as active for each respective day.

### Freezing and activity correlations

For correlating neuronal activity and mouse freezing behavior (Figure 3J-K), we calculated the mean value of the freezing and activity trace for the 20-second time window from 10 seconds before to 10 seconds after cue offset then averaged for each mouse for each day. We then calculated the Pearson correlation for each group for each day with a 95% CI.

### Tuning curve correlations

For analyzing differences in representational drift, we first smoothed activity traces using a 0.5 second moving average kernel, then used averaging to downsample the traces to 0.5Hz. We then calculated the tuning curve correlations between the activity traces of individual cells during the first 7 seconds of different cue presentations. We then averaged across the recorded neuronal population for each mouse.

### Hierarchical clustering

Calcium traces were averaged across trials for each recording day, to obtain a single mean trace per neuron. Traces were smoothed using a 1-second moving average to reduce noise before clustering. Neurons were concatenated into one pooled matrix where every row is the activity of one neuron from one mouse on one day. We then performed hierarchical clustering via pairwise correlation distances for each neuron in the pooled matrix. Cluster membership was defined by clipping the dendrogram at 4 clusters to preserve major branching structure while avoiding over-segmentation (Figure 4E). Cluster identities were then mapped back to each neuron’s original mouse & day for all subsequent analysis.

To obtain cluster stability across days (Figure 4G, supplemental Fig 6), we used the percent of neurons assigned to cluster c on day A that remained in cluster c on day B, a conditional probability of P(c_dayB | c_dayA). We only used the subset of neurons that were detected in both days A and B.

To obtain temporal correlations per cluster across days, activity from all neurons belonging to a given cluster was averaged within mouse for each cue presentation to generate a cluster population activity vector. Pairwise Pearson correlations were computed between cluster population vectors across cues and days. Self-correlations were excluded. Correlations were then grouped according to the number of days between cue pairs.

## Disclosures

A.P.K, C.P., J.H.K, and A.F.R are co-inventors on a patent application related to psychedelics but not related to this work. Dr. Kaye previously received contracted research funding from Transcend Therapeutics. Drs. Kaye, Jefferson and Pittenger previously received research funding from Freedom Biosciences. In the past three years, Dr. Pittenger has consulted for Biohaven Pharmaceuticals, Freedom Biosciences, Transcend Therapeutics, UCB BioPharma, Mind Therapeutics, Ceruvia Biosciences, F-Prime Capital Partners, and Madison Avenue Partners; has received research support from Biohaven Pharmaceuticals, Freedom Biosciences, and Transcend Therapeutics; owns equity in Alco Therapeutics, Mind Therapeutics, and Lucid/Care; receives royalties from Oxford University Press and UpToDate; and holds pending patents on pathogenic antibodies in pediatric OCD and on novel mechanisms of psychedelic drugs. Dr. Krystal has served as a consultant for Aptinyx, Inc.; Biogen, Idec, MA; Bionomics, Limited (Australia); Boehringer Ingelheim International; Clearmind Medicine, Inc.; Cybin IRL (Ireland Limited Company); Enveric Biosciences; Epiodyne, Inc.; EpiVario, Inc.; Janssen Research & Development; Jazz Pharmaceuticals, Inc.; Otsuka America Pharmaceutical, Inc.; Perception Neuroscience, Inc.; Praxis Precision Medicines, Inc.; Spring Care, Inc.; Sunovion Pharmaceuticals, Inc. Dr. Krystal has served as a scientific advisory board member for: Biohaven Pharmaceuticals; BioXcel Therapeutics, Inc. (Clinical Advisory Board); Cerevel Therapeutics, LLC; Delix Therapeutics, Inc.; Eisai, Inc.; EpiVario, Inc.; Freedom Biosciences, Inc.; Jazz Pharmaceuticals, Inc.; Neumora Therapeutics, Inc.; Neurocrine Biosciences, Inc.; Novartis Pharmaceuticals Corporation; Praxis Precision Medicines, Inc.; PsychoGenics, Inc.; Tempero Bio, Inc.; Terran Biosciences, Inc. In the past 3 years, Dr. Krystal is an inventor on patents licensed by Yale University to Janssen Pharmaceuticals, Biohaven Pharmaceuticals, Spring Health, Freedom Biosciences, and Novartis Pharmaceuticals. The remaining authors declare no conflict of interest.

## Supplemental figures

**Sup fig 1.**
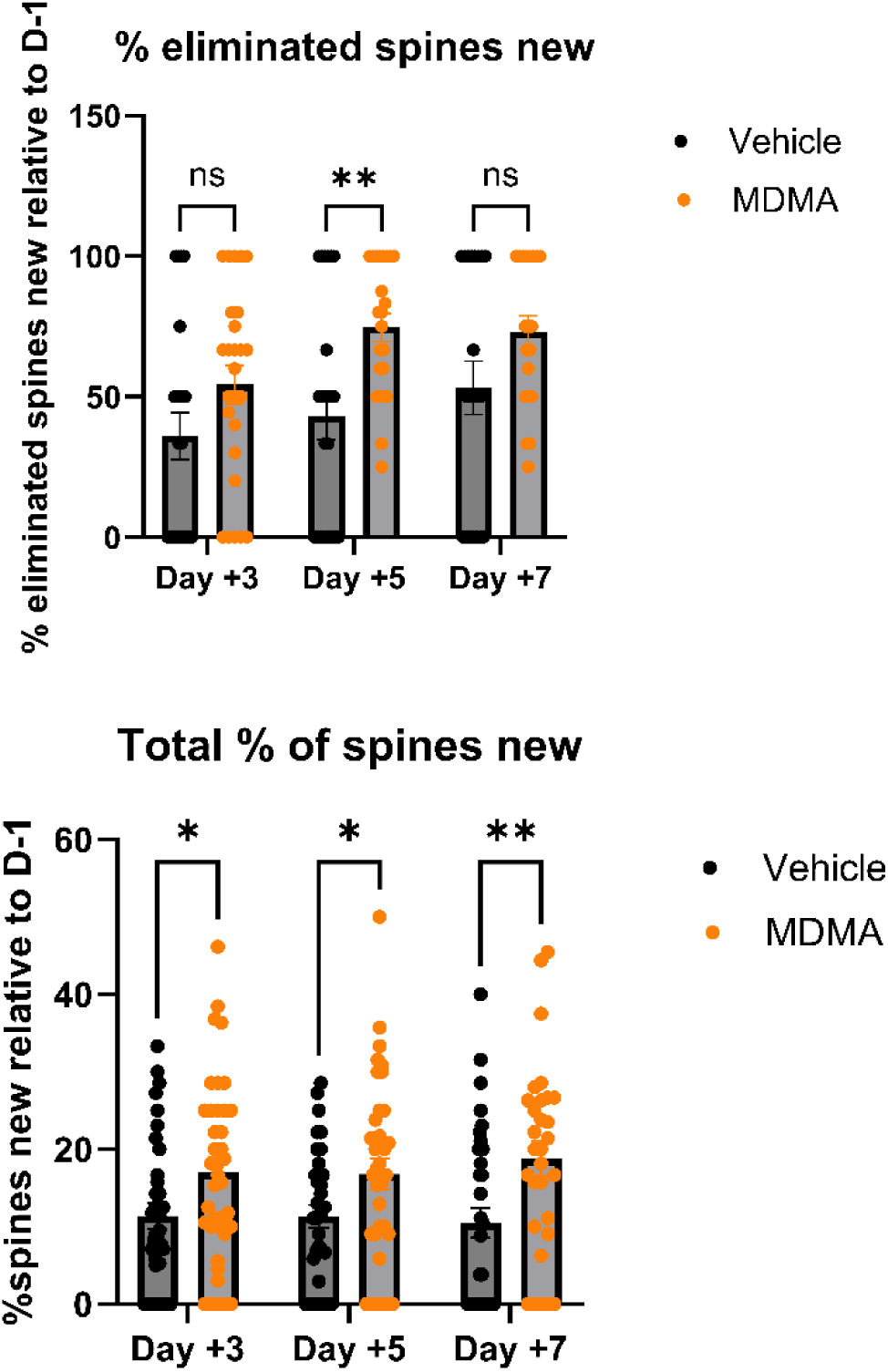

**Sup Fig 2.**
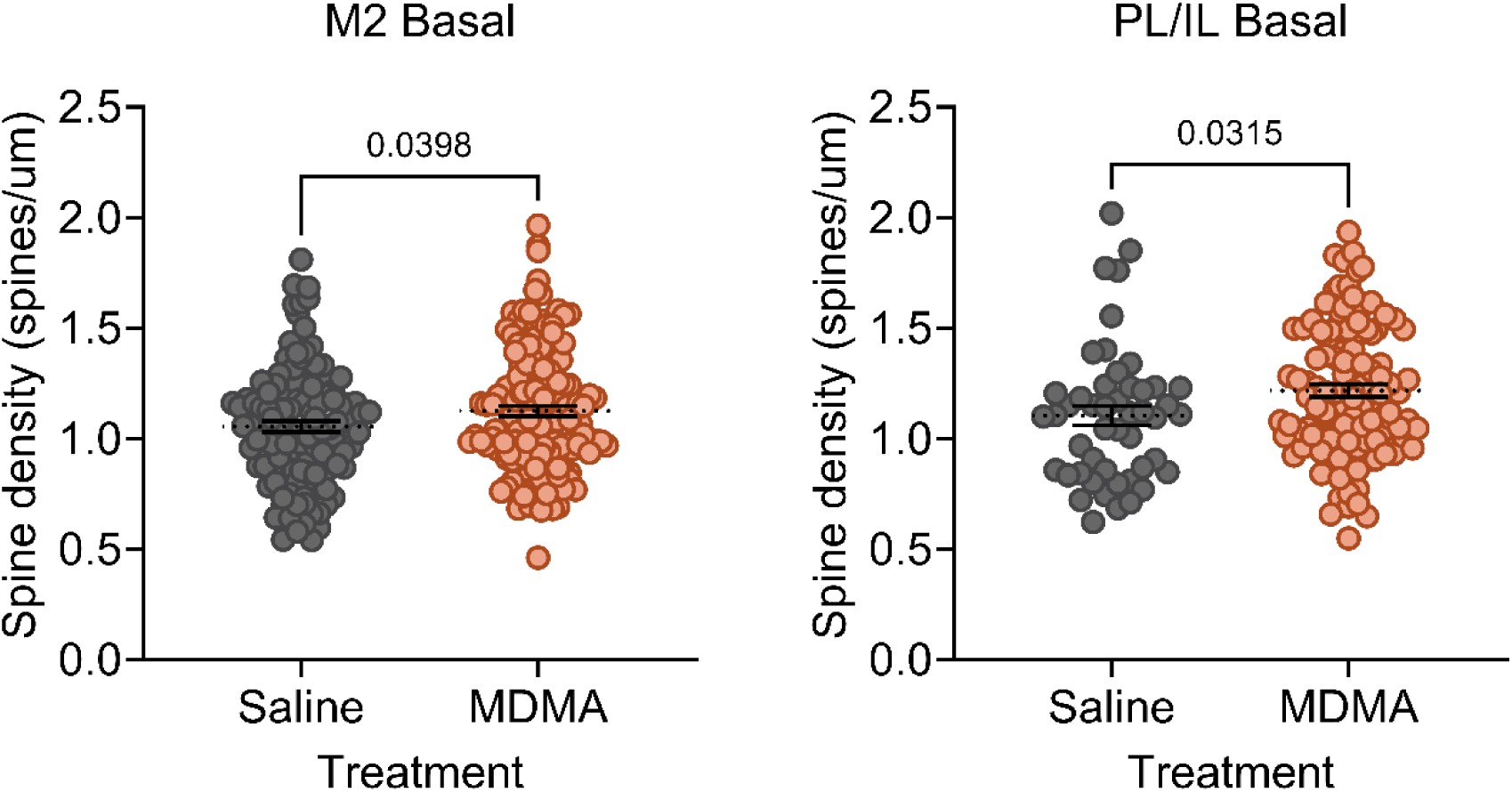
– MDMA increases spinogenesis in basal dendrites in mPFC. Quantification of spine density in basal dendrites from Cg1/M2 (A) and PL/IL (B) 24 hours after saline or MDMA injection. Two-tailed *t*-test: p = 0.0398 (A), p = 0.0315 (B).

**Supplemental Fig 3.**
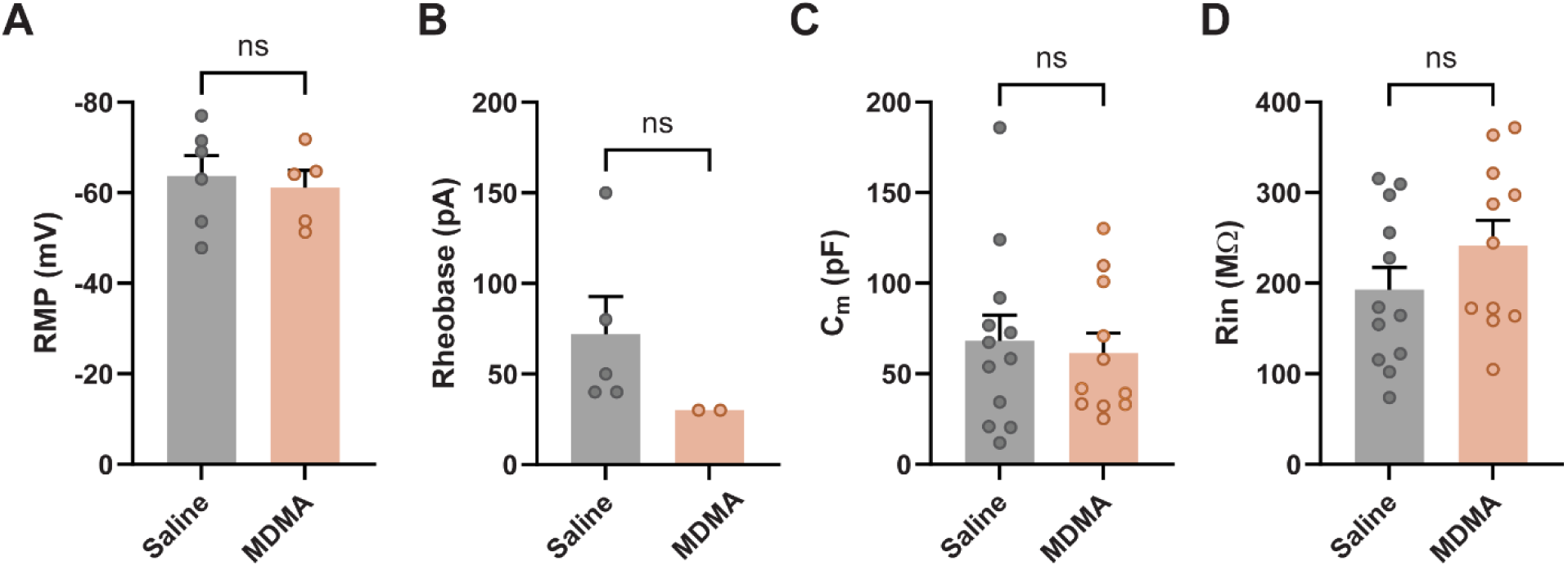
– Intrinsic membrane properties of IL L5 pyramidal neurons are not altered by MDMA. (A) Resting membrane potential (RMP), (B) rheobase, (C) membrane capacitance (Cₘ), and (D) input resistance (Rᵢₙ) of IL layer 5 pyramidal neurons 24 hours after injection of saline (gray) or MDMA 7.8 mg/kg (orange). No significant differences were observed between treatment groups for any measure (Welch’s t-tests, all p > 0.05). Data are plotted as mean ± SEM with individual neurons shown.

**Supplemental Fig 4.**
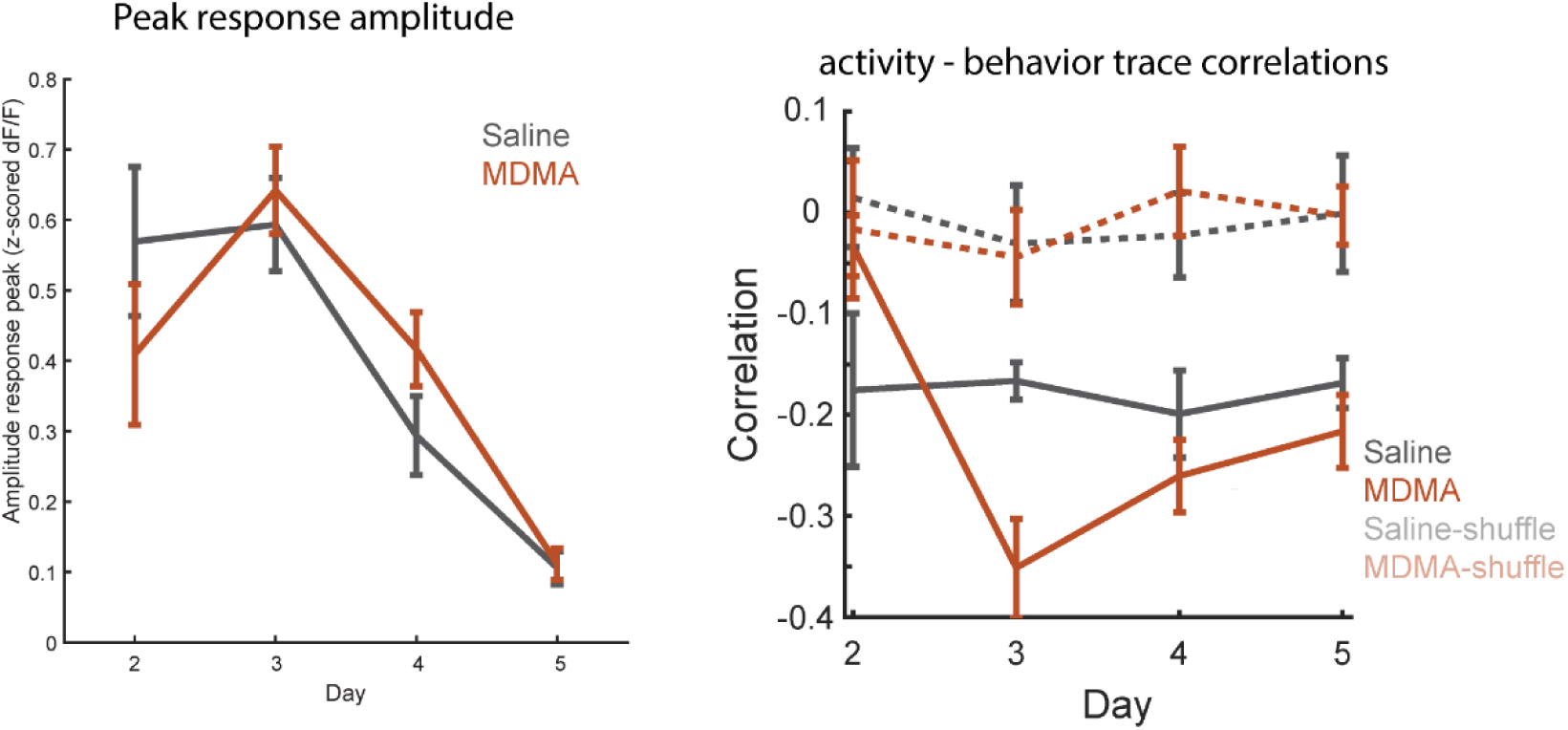
- Peak cue-evoked response amplitude and activity–behavior trace correlations across extinction days. (A) Peak amplitude of the population Ca²⁺ response (z-scored ΔF/F) to the auditory cue across extinction days for saline-(gray) and MDMA-(orange) treated mice. Both groups show similar peak response magnitudes that decline over extinction training. Data plotted as mean ± SEM. (B) Correlation of Ca2+ activity (Fig 1H) and freezing (Fig 1F) temporal traces, shows a sustained negative correlation in MDMA-treated mice. Solid lines show observed correlations for saline (gray) and MDMA (orange) groups; dashed lines show shuffle controls in which trial identity was randomly permuted. MDMA-treated mice showed a stronger negative correlation between IL activity and freezing over extinction testing days (days 3–5), while saline-treated mice maintain a relatively stable correlation. Data plotted as mean ± SEM.

**Supplemental Fig 5.**
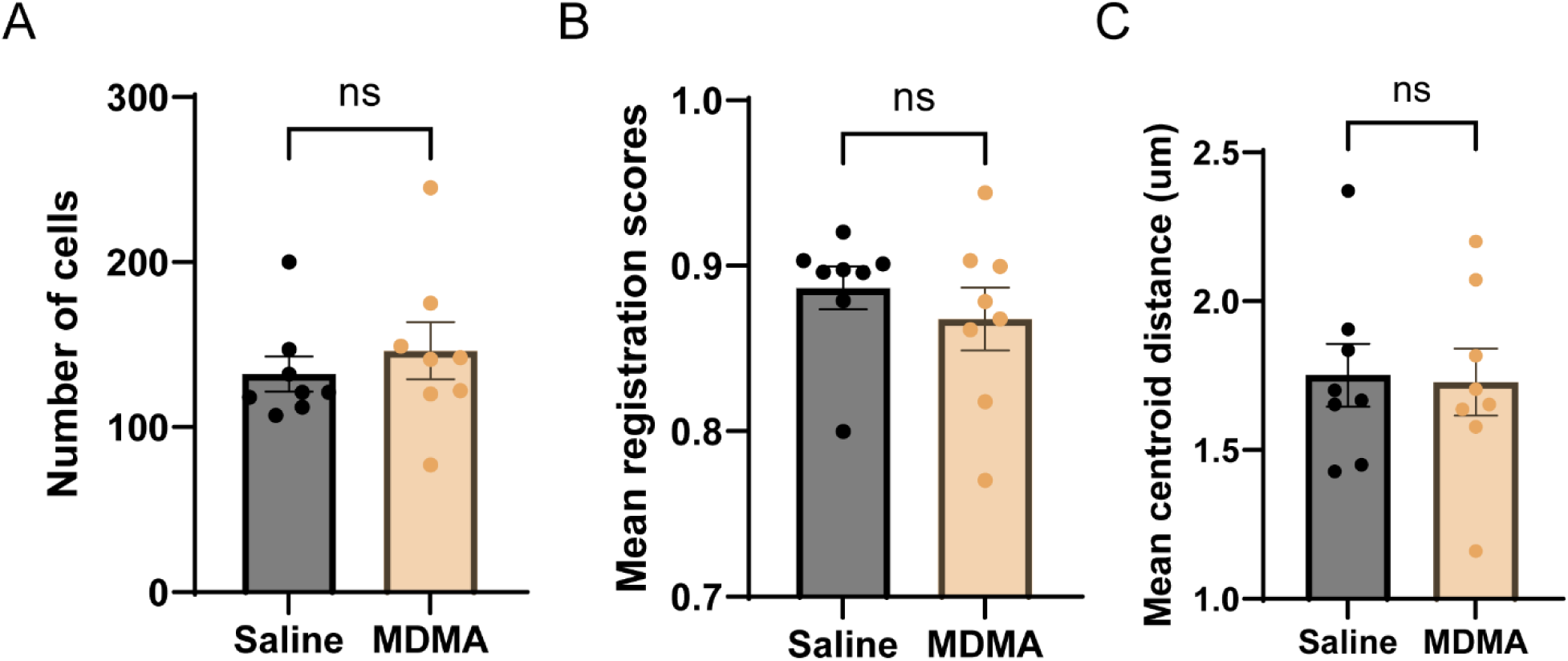
– Registration quality is similar across treatment groups. number of registered cells, mean registration score per cell, and mean centroid distances were similar for saline and MDMA-treated mice (p= 0.5, 0.43, and 0.88, respectively)

**Sup Fig 6.**
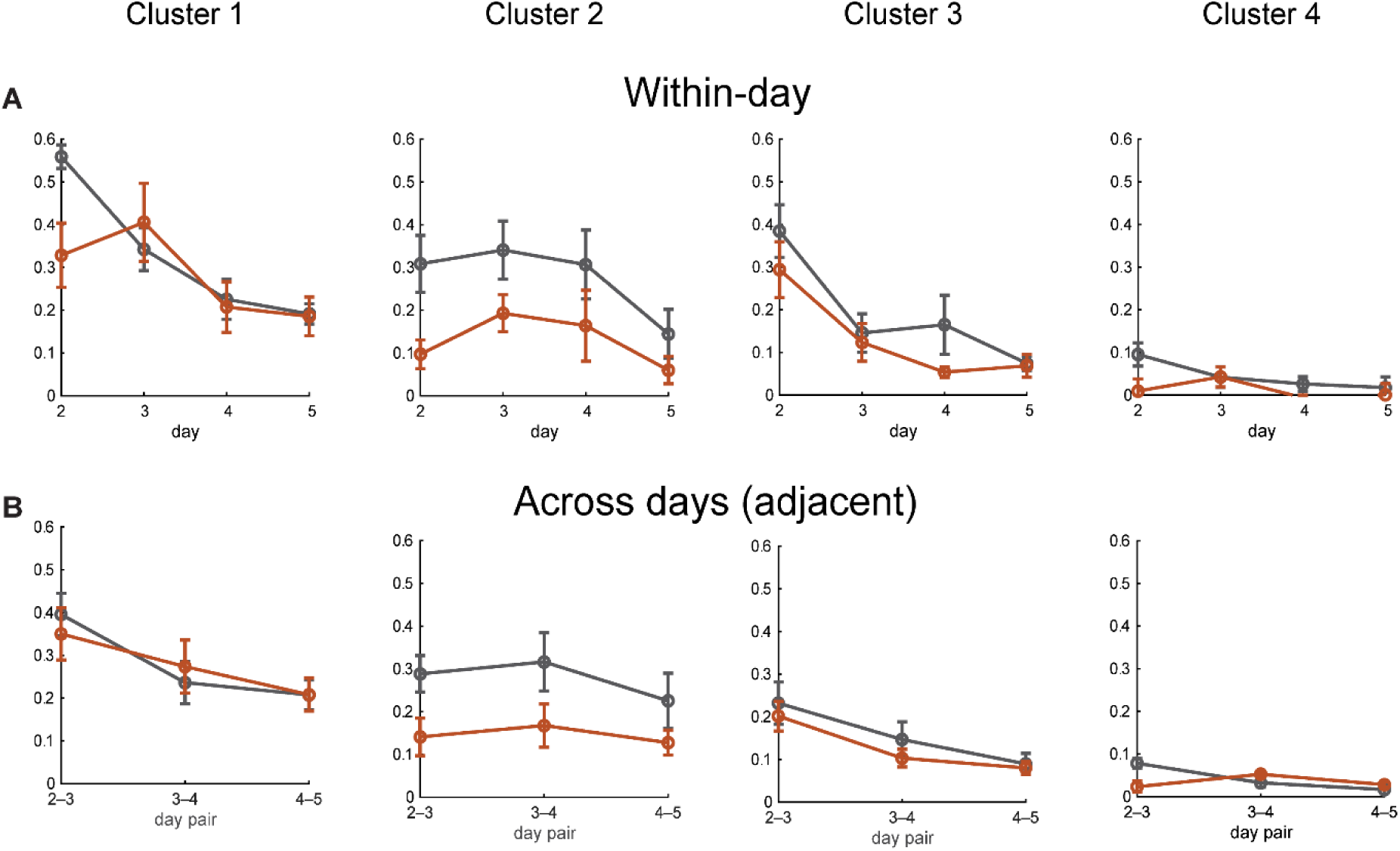
– dynamics of cue-evoked tuning correlations for cells within each functional cluster. Pairwise correlations of individual cells’ tuning curves during entire cue presentation for cells identified in each of the four functional clusters, shown for within-day comparisons (A), and across adjacent day pairs (B). Gray = saline, orange = MDMA. Data plotted as mean ± SEM. **Cluster 1**: Within-day correlations showed a significant main effect of treatment (F(1,48) = 9.76, p = 0.003), day (F(3,48) = 14.27, p < 0.001), and a day × cohort interaction (F(3,48) = 4.28, p = 0.009). No significant cohort effects were found for adjacent day-pairs (p > 0.16). **Cluster 2**: Within-day correlations showed a significant main effect of treatment (F(1,48) = 7.70, p = 0.008), with MDMA-treated mice exhibiting lower correlations, but no day × cohort interaction (p = 0.72). The treatment effect was also significant for adjacent day pairs (F(1,29) = 4.89, p = 0.035). **Cluster 3**: No significant treatment effects were observed (p > 0.15). Within-day correlations decreased over extinction days (F(3,48) = 9.51, p < 0.001), and adjacent day-pair correlations also declined (F(2,29) = 6.03, p = 0.006), indicating time-dependent decorrelation that was shared across treatment groups. **Cluster 4**: Within-day correlations showed a significant main effect of treatment (F(1,47) = 7.20, p = 0.010). Adjacent day-pair analysis revealed significant effects of treatment (F(1,27) = 13.09, p = 0.001) and a pair × treatment interaction (F(2,27) = 7.79, p = 0.002).

## Notes

### Competing Interest Statement

The authors have declared no competing interest.

### Funding

- United States Department of Defense: HT9425-23-1-0458
- National Institutes of Health: R56 MH137464

